# Non-alcoholic components of Pelinkovac, a Croatian wormwood-based strong liquor, counteract the inhibitory effect of high ethanol concentration on catalase *in vitro*

**DOI:** 10.1101/2022.01.07.475357

**Authors:** Jan Homolak, Ana Babic Perhoc, Mihovil Joja, Ivan Kodvanj, Karlo Toljan, Davor Virag

## Abstract

Antioxidant enzyme catalase protects the cells against alcohol-induced oxidative stress by scavenging free radicals and metabolizing alcohol. Concentrations of ethanol present in alcoholic beverages can inhibit catalase and foster oxidative stress and alcohol-induced injury. Non-alcoholic components of pelinkovac counteract the inhibitory effects of high ethanol concentration and acidic pH on catalase *in vitro*.

## Introduction

Although there is evidence supporting beneficial effects of low levels of alcohol consumption in some conditions (e.g. cardiovascular disease, diabetes) its use has been linked to myriad acute and chronic diseases and has been recognized as a leading risk factor for death and disability worldwide (Griswold et al., 2018). Elucidating molecular mechanisms responsible for the detrimental consequences of excessive alcohol consumption might provide the foundations for the development of preventive and therapeutic strategies to reduce the alcohol-attributable burden of disease.

Ethanol metabolism produces a substantial amount of reactive oxygen species (ROS) and oxidative stress has been proposed as a pathophysiological perpetrator responsible for the harmful effects of excessive alcohol consumption (Albano, 2006; Yue et al., 2021). Catalase (CAT; EC 1.11.1.6), plays a dual role in pathophysiological processes associated with alcohol consumption. Being the main regulator of the endogenous ROS H_2_O_2_ (Góth et al., 2004), CAT protects the cell against alcohol-induced oxidative stress by removing excess H_2_O_2_. Furthermore, CAT is directly involved in alcohol metabolism by catalyzing the reduction of H_2_O_2_ to H_2_O in the process in which ethanol (used as an electron donor) is oxidized to acetaldehyde (Comporti et al., 2010). Although CAT-mediated alcohol metabolism acts as a secondary metabolic pathway in the liver under „normal circumstances“, it seems to be a predominant pathway in some conditions (e.g. during fasting) and in some organs (e.g. in the brain where alcohol dehydrogenase is not present)(Yue et al., 2021). CAT expression is reduced in patients with severe alcoholic hepatitis (Yue et al., 2021), and the CAT gene seems to be associated with susceptibility to alcohol dependence (Plemenitas et al., 2015). Interestingly, it has been recently shown that CAT-deficient mice are more susceptible to alcohol-induced inflammation and oxidative stress and that the switch from the ROS-generating CYP2E1 pathway to ROS-scavenging CAT-mediated metabolism of ethanol can reverse alcohol-induced liver damage in mice (Yue et al., 2021).

Apart from being a scavenger of ethanol metabolism-generated ROS and an important metabolic pathway for alcohol detoxification, CAT may also be directly involved in the etiopathogenesis of alcohol toxicity. It has been shown that wine concentrations of ethanol are sufficient for a substantial inhibition of CAT via a non-competitive mechanism that involves binding to the compound I form of the enzyme (Temple and Ough, 1975), and oxidative stress in rats exposed to alcohol has been associated with reduced CAT activity (Das and Vasudevan, 2005; Jurczuk et al., 2004).

Ethanol and water are the main constituents of alcoholic beverages, however, most also contain a substantial number of non-alcoholic components (e.g. tannins and polyphenols) which have been proposed to mediate at least some of the beneficial health effects of moderate alcohol consumption (Arranz et al., 2012). Many non-alcoholic components of alcoholic beverages have shown antioxidant effects *in vitro* and *in vivo* (Rodrigo et al., 2011) including potentiation of CAT activity (e.g. (Noguer et al., 2012)).

In the present study, we wanted to assess the effects of pelinkovac on the activity of CAT. Pelinkovac is a traditional Croatian strong liquor based on wormwood, which is believed, according to folk tales, to have healing properties. We hypothesized that pelinkovac would be able to inhibit CAT due to the high content of ethanol (~30%), but that non-alcoholic components of pelinkovac from the macerate of the aromatic herbs may be able to counteract some of the effects of ethanol.

The aim was to determine whether: i) pelinkovac can inhibit CAT *in vitro;* ii) two different types of pelinkovac (Antique and Epulon) exert different effects on CAT; iii) selected aromatic herbs present in pelinkovac potentiate or counteract the effects of ethanol and pH on CAT.

## Materials and Methods

### Sample preparation

Antique Pelinkovac (A) was obtained from Badel (Zagreb, Croatia) and Epulon Pelinkovac (E) was obtained from Rossi (Vizinada, Istria, Croatia). pH-adjusted Epulon (pHE) (introduced to assess whether potential differences of the effects of A and E on CAT might be due to pH) was prepared by adjusting the pH of E (4.28) to the original pH value of A (3.82) with HCl. Ethanol-based control solutions for A (ETA) and E (ETE) (used to assess whether the observed effects on CAT were due to ethanol in combination with low pH) were prepared by adjusting the pH of i) 35% (v/v) ethanol in ddH_2_O to 3.82 (ETA), and ii) 30% (v/v) ethanol in ddH_2_O to 3.82 (ETE). CAT solution was prepared by dissolving 1 mg of lyophilized powder of CAT isolated from bovine liver (Sigma-Aldrich, USA) in 10 ml of phosphate-buffered saline (PBS)(pH 7.4).

### Catalase activity

CAT activity was assessed using an adaptation of the method first introduced by Hadwan (Hadwan, 2018) as described and validated in (Homolak, 2021; Homolak et al., 2022). In brief, 20 μL of CAT solution was placed in a well with 5 μL ddH_2_O (CAT), A (CAT+A), E (CAT+E), pHE (CAT+pHE), ETA (CAT+ETA), or ETE (CAT+ETE). For baseline absorbance (t_0_ = 0 s), 150 μL of the Co(NO_3_)_2_ stop solution (5 mL Co(NO_3_)_2_ x 6 H_2_O (0.2 g in 10 mL ddH_2_O) + 5 mL (NaPO_3_)_6_ (0.1 g in 10 mL ddH_2_O) added to 90 mL of NaHCO_3_ (9 g in 100 mL ddH_2_O)) was first added to samples followed by 50 μL of 2,4,6,8, and 10 mM H_2_O_2_ in PBS. For the activity measurements, samples were first incubated with 50 μL of 2,4,6,8, and 10 mM H_2_O_2_ in PBS, and the reaction was stopped with 150 μL of the Co(NO_3_)_2_ stop solution after 15, 30, 60, and 120 s. The concentration of H_2_O_2_ was determined indirectly by measuring the absorbance of the carbonato-cobaltate (III) complex ([Co(CO_3_)_3_]Co) at 450 nm using the Infinite F200 PRO multimodal microplate reader (Tecan, Switzerland), ti = 15 s was used for the calculation of the maximum velocity at a saturated concentration (V_max_) and Michaelis-Menten constant (K_M_). All measurements were done at 25 °C.

### Data analysis

Data were analyzed in R (4.1.0). Hadwan’s method (Hadwan, 2018) uses independent cross-sectional observations of H_2_O_2_ concentration to estimate CAT activity. Consequently, total enzyme activity at t_2_ = 120 s and enzyme activity at different substrate concentrations at t_1_ = 15 s was based on permutation-derived estimates. The effects of pelinkovac and control solutions on CAT were analyzed using linear regression. Model assumptions were checked using visual inspection of residual and fitted value plots and model outputs were reported as point estimates of group differences with 95% confidence intervals. V_max_ and K_M_ were estimated from the double reciprocal Lineweaver-Burk plots and using the Levenberg-Marquardt non-linear least-squares algorithm (Elzhov et al., 2016)(Supplement).

## Results

Both A and E exerted an inhibitory effect on CAT activity (Fig 1A,B). The H_2_O_2_ decomposition rate was reduced on average by 1.15 and 0.87 mM in the presence of A and E respectively (Fig 1A,B). ETA and ETE exerted an even more pronounced inhibitory effect suggesting that aromatic herbs present in A and E „protect” CAT against the inhibitory effect of pH and alcohol. Interestingly, the difference between the inhibitory effect of A and ETA (δH_2_O_2_=1.28 mM) was greater than the difference between E and ETE (δH_2_O_2_=0.45 mM) indicating a more pronounced protective“ effect of aromatic herbs in A. pHE and ETE produced a comparable level of inhibition that was less pronounced than that observed with ETA implying that even relatively low increase in ethanol concentration can significantly potentiate the inhibition of CAT.

**Fig 1.**
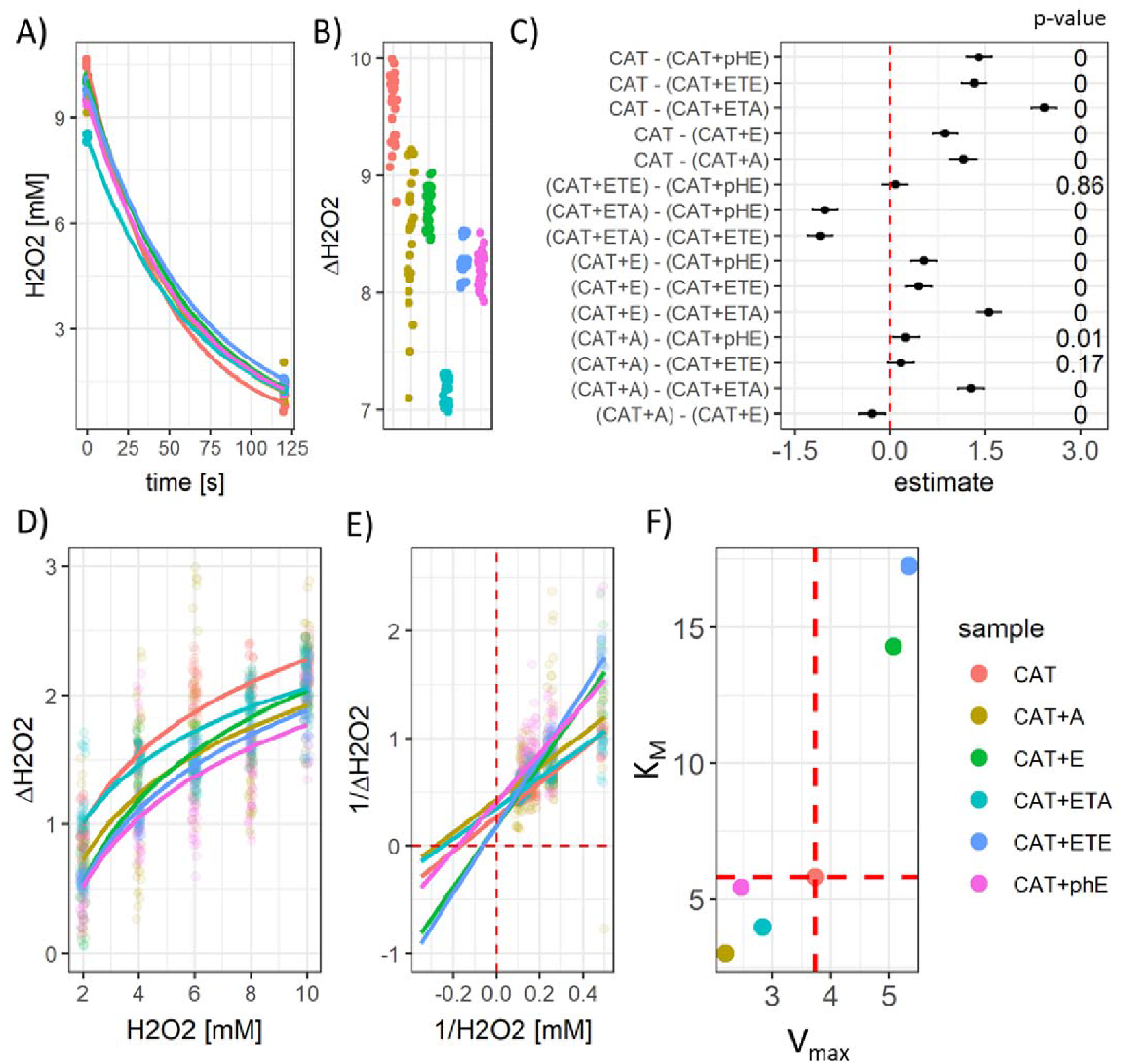
The effect of pelinkovac on catalase (CAT) *in vitro.* A) The effect of Antique Pelinkovac (A), Epulon Pelinkovac (E), and respective control solutions - pH-adjusted Epulon (pHE), dummy ethanol-based control solutions for A (ETA) and E (ETE), on the activity of CAT. B) The effect of pelinkovac and control solutions on total CAT activity in 120 s. C) Model output represented as point estimates of condition differences with 95% confidence intervals. D) The Michaelis-Menten plot of CAT activity at different substrate concentrations. E) The Lineweaver-Burk double reciprocal plot with the reciprocal velocity as the dependent variable and the reciprocal of substrate concentration as the independent variable. F) Visual representation of the maximum velocity at a saturated concentration (V_max_) and Michaelis-Menten constant (K_M_) for each experimental condition derived from E).

Enzyme kinetics analysis revealed that A and ETA primarily inhibit CAT by binding to the enzymesubstrate complex as evident from the uncompetitive inhibition pattern characterized by reduced V_max_ and K_M_ (Fig 1 D-F). Interestingly, the effect of pHE resembled noncompetitive inhibition, and the effects of E and ETE were accompanied by a perplexing decrement of substrate binding affinity with increased catalytic power of CAT (Fig 1 D-F).

## Discussion

The presented results demonstrate that pelinkovac can inhibit CAT *in vitro.* Although this was expected due to high alcohol concentration it was surprising to see that pelinkovac (A, E) inhibited CAT to a lesser extent than pH-adjusted control solutions of ethanol (ETA, ETE) suggesting that nonalcoholic components of pelinkovac may counteract the effects of ethanol on CAT. The exact composition of pelinkovac is a trade secret, but it is known that the production process involves alcohol maceration of „selected aromatic herbs“, macerate aging, and distillation in copper (“Antique Pelinkovac,” 2022). The main herbal constituent of pelinkovac is wormwood *(Artemisia absinthium* L.) – a well-known species of the genus *Artemisia* recognized as a medicinal herb and declared *„the most important master against all exhaustions“* in medieval Europe due to its many favorable biological effects (Szopa et al., 2020). Although the direct effect of wormwood on CAT has never been reported to the best of our knowledge, the herb has shown antioxidant effects (Szopa et al., 2020) indicating some of its many bioactive compounds may also exert a modulatory effect on CAT. Although the selection of other herbs depends on the brand of pelinkovac, usually around 20 are present many of which belong to the class of medicinal herbs (e.g. *Angelica archangelica* L., *Pimpinella anisum* L., *Vaccinium* L., *Juniperus communis* L., *Acorus calamus* L., *Teucrium montanum* L., *Hyssopus officinalis* L., *Fragaria vesca* L., *Salvia officinalis* L., *Centaurium erythraea* subsp. *erythraea, Foeniculum vulgare* Mill., *Coriandrum sativum* L., *Thymus serpyllum* L., *Melissa officinalis* L., *Origanum majorana* L., *Mentha × piperita* L., *Calendula officinalis* L., *Inula helenium* L., *Salvia rosmarinus* Spenn.). The overview of the antioxidant effects and the protective effect on CAT of the aforementioned herbs is beyond the scope of this manuscript, however many have shown protective effects on CAT indicating bioactive compounds from these herbs may also contribute to the observed protective effect against ethanol-induced inhibition of the enzyme *in vitro.*

Although the *in vitro* effects of pelinkovac reported here cannot be directly translated to the repercussions for the CAT activity *in vivo,* it can be assumed that pelinkovac may exert its inhibitory action in the oral cavity, and on the intraluminal and mucosal gastrointestinal CAT of both human and bacterial origin which raises many questions that may be relevant for human health. In this experiment, a substantial level of CAT inhibition was achieved regardless of all tested solutions being diluted to only 6.6% of their original concentration in the working buffer indicating that even a considerable dilution in saliva and gastric juice would not be able to mitigate the observed effect. A possible effect on the bacterial CAT may mediate the effect of pelinkovac (and other alcoholic beverages) on the qualitative and quantitative features of microbiota shifting the multispecies microbial community towards dysbiosis or eubiosis as both H_2_O_2_ and CAT are involved in the commensal-pathobiont crosstalk (e.g. (Herrero et al., 2018)). The effect on mucosal human CAT may also be relevant as excess H_2_O_2_ and suppressed CAT have been reported in pathological conditions such as inflammatory bowel disease and gastric adenocarcinoma, and potentiating intestinal CAT activity in the gut is being explored as a potential therapeutic option in these conditions (Alothman et al., 2021).

Based on the aforementioned, understanding the effects of other alcoholic beverages other than pelinkovac on CAT activity *in vitro* and assessment of the effect of alcoholic beverages in general on the CAT *in vivo* may be important for elucidating the pathomechanisms responsible for the alcohol-attributable burden of disease. Apart from that, future research should explore the observed protective effect of non-alcoholic components of pelinkovac for their protective effects on CAT.

## Conclusion

A famous Croatian wormwood-based liquor pelinkovac inhibits CAT *in vitro* probably due to high ethanol content. Pelinkovac inhibits CAT to a lesser extent than pH and ethanol-content matched control solutions indicating some non-alcoholic components of pelinkovac counteract the inhibitory effect of ethanol. Future research should explore whether the observed effect is relevant for oral and gastrointestinal intraluminal and mucosal CAT of human and microbial origin, and clarify which pelinkovac-derived compounds are responsible for mitigating the inhibitory effects of high ethanol concentration on CAT.

## Data availability statement

Raw data can be obtained from the corresponding author’s GitHub: https://github.com/janhomolak

## Author’s contributions

**JH** conceived and conducted the experiment and wrote the first draft of the manuscript. **ABP** obtained pelinkovac for the defense of her doctoral thesis. **ABP, MJ, IK, KT,** and **DV** critically contributed to the discussion of the pataphysiological aspects of pelinkovac and to the revision of the manuscript. All authors read and approved the final version of the manuscript.

